# Epigenetic Germline Variants Predict Cancer Prognosis and Risk and Distribute Uniquely in Topologically Associating Domains

**DOI:** 10.1101/2023.07.04.547722

**Authors:** Shervin Goudarzi, Meghana Pagadala, Adam Klie, James V. Talwar, Hannah Carter

## Abstract

Cancer is a highly heterogeneous disease caused by genetic and epigenetic alterations in normal cells. A recent study uncovered methylation quantitative trait loci (meQTLs) associated with different levels of local DNA methylation in cancers. Here, we investigated whether the distribution of cancer meQTLs reflected functional organization of the genome in the form of chromatin topologically associated domains (TADs), and evaluated whether cancer meQTLs near known driver genes have the potential to influence cancer risk or progression. At TAD boundaries, we observed differences in the distribution of meQTLs when one or both of the adjacent TADs was transcriptionally active, with higher densities near inactive TADs. Furthermore, we found differences in cancer meQTL distributions in active versus inactive TADs and observed an enrichment of meQTLs in active TADs near tumor suppressors, whereas there was a depletion of such meQTLs near oncogenes. Several meQTLs were associated with cancer risk in the UKBioBank, and we were able to reproduce breast cancer risk associations in the DRIVE cohort. Survival analysis in TCGA implicated a number of meQTLs in 13 tumor types. In 10 of these, polygenic meQTL scores were associated with increased hazard in a CoxPH analysis. Risk and survival-associated meQTLs tended to affect cancer genes involved in DNA damage repair and cellular adhesion and reproduced cancer-specific associations reported in prior literature. In summary, this study provides evidence that genetic variants that influence local DNA methylation are affected by chromatin structure and can impact tumor evolution.

## Introduction

Cancer is a heterogeneous disease and common treatments like chemotherapy have only a 55% response rate^1^. Precision medicine and biomarker analysis can tailor treatment options and optimize outcomes. Genetic factors, such as germline and somatic mutations, contribute to heterogeneous disease risk and progression. For example, germline variants in the *BRCA2* gene can greatly increase the risk of developing breast and ovarian cancer^2^. Epigenetic factors including DNA methylation, histone modification, and acetylation also play a key role in cancer progression. Recently, promising therapeutics have been developed that inhibit DNA Methyltransferases (DNMTs), reducing tumor growth in breast cancer and highlighting the importance of DNA methylation and other epigenetic factors in carcinogenesis^2, 3^. However, the interplay between epigenetics and genetics in cancer risk and progression remains mostly elusive.

Methylation Quantitative Trait Loci, or meQTLs, are SNPs that significantly correlate with DNA methylation at CpG sites. These SNPs provide a bridge between genetic variation and corresponding epigenetic effects shown to correlate with cancer risk^4^. Disruptions in DNA methylation are well-known in the context of cancer; DNA is frequently hypermethylated at promoter regions of tumor suppressor genes while hypomethylated at the promoters of oncogenes, and there is an inverse correlation with gene expression^53^. Promoter hyper-and hypo-methylation has been of specific interest due to its role in regulating the expression of cancer genes including suppression of tumor suppressor genes like BRCA^6^ and the expression of oncogenes like L1NE1^7^. Subsequently, germline SNPs that acted as meQTLs were shown to predict risk in many cancer types like breast and lung, regulating expression and methylation of genes like FBXO-18^4^.

The organization of the genome into 3D structures may further modify the potential of genetic variants to interact with epigenetic factors in a disease specific manner^8^. Topologically Associating Domains (TADs), are isolated regions of highly-interacting and folded chromatin separated by insulator proteins. TADs are important for maintaining controlled patterns of local gene regulation and provide a framework for transcriptionally similar genes and SNPs to interact with one another^9^. In fact, because TADs have been found to be highly stable across tissue types, they provide valuable context for understanding the genome’s functional landscape allowing the study of genetic variation in the context of 3D chromatin structure ^10^. Mutational burden of somatic mutations within the context of cancer demonstrated correlation with TADs ^11^. In addition, genes within TADs demonstrate correlated gene expression and histone modification ^12, 13^, allowing us to group similar acting genes and SNPs, narrowing a search for potentially cancer related SNPs.

In this study, we integrate genetic correlates of DNA methylation across 23 cancer types (i.e. cancer meQTLs) and TAD domains to better understand how 3-D chromatin structure might determine the potential of meQTLs to influence cancer risk and survival. We focus on meQTLs near TADs containing key cancer-related genes. Analyzing the location and distribution of such variants across the genome, we find that methylation-related germline variants, or meQTLs, in cancer do not lie uniformly across the genome and the occurrence of TAD boundaries correlates with significant cancer meQTL presence. In addition, meQTLs closely related to cancer progression show specific nonrandom distribution in TAD domains. Then we assessed whether meQTLs near cancer genes could predict cancer survival and risk and found significant prediction power of these meQTLs across multiple cancer types. Our study suggests that the potential of meQTLs to contribute to cancer risk and progression depends in part on local genome architecture and chromatin state.

## Results

### Active TADs are associated with less DNA methylation at cancer meQTLs

We identified 1100 TADs shared across 5 cell lines (GM12878, HMEC, HUVEC, IMR90, and NHEK) and categorized them into “Mixed”, “Inactive-1”, “Inactive-2”, “Active-1”, and “Active-2” groups using chromatin state information (**Figure 1A**). Combining the active and inactive groups resulted in 222 active, 626 inactive and 252 mixed TADs. DNA methylation is linked with TAD activity via nucleosome positioning and chromatin condensation ^14^ as well as to regulation of gene expression, where promoter CpG methylation is associated with gene silencing ^15^. We compared our categorization of TAD activity with genome-wide DNA methylation in promoter regions defined based on the ENCODE Screen Pipeline. Promoters in active TADs showed overall lower levels of methylation whereas those in inactive TADs had a higher level of methylation (Kruskal-Wallis, p-value < 0.001) (**Figure 1B**), supporting that promoter methylation silencing aligns with categorization of TADs into transcriptionally different groups, namely into “active” and “inactive”.

**Figure 1.**
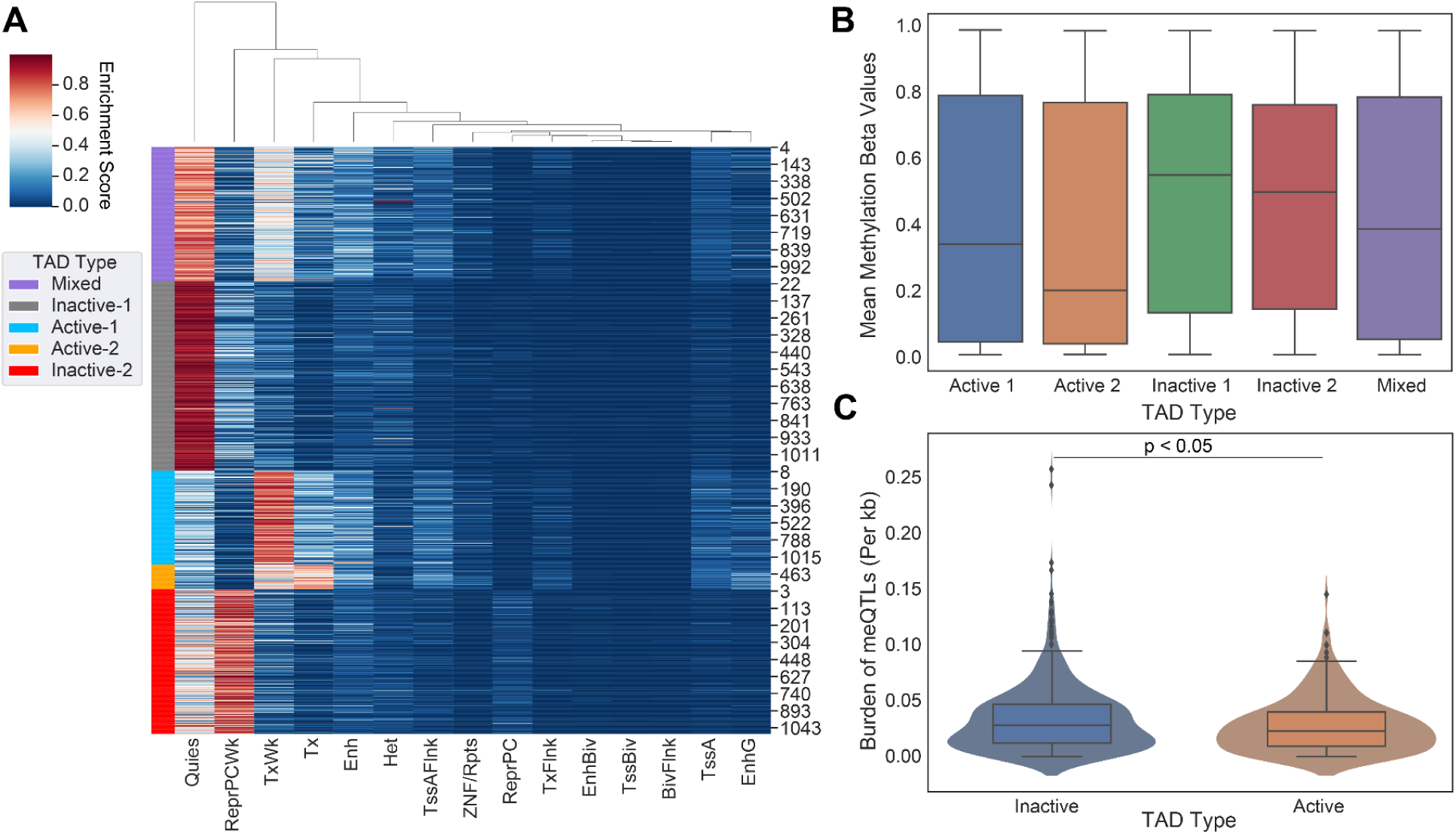
Evaluating DNA methylation and meQTL burden in Topologically Associated Domains (TADs). (**A**) 5 state-based K-Means clustering of common TAD domains (n=1100) between 5 human cell lines (GM12878, HMEC, HUVEC, IMR90, and NHEK). Purple indicates TADs classified as a “Mixed”, Gray as “Inactive-1”, Light Blue as “Active-1”, Orange as “Active-2”, and Red as “Inactive-2”. Combining active and inactive categories leads to 222 Active, 626 Inactive, and 252 Mixed TADs. (**B**) On average, inactive TADs have higher DNA methylation levels than active TADs (*p*-value< 0.001). These results are supported by previous literature concerning promoter methylation and transcriptional activity. (**C**) Number of meQTLs across inactive TADs versus active TADs are shown. meQTL counts per TAD were normalized by TAD length in base pairs. Active TADs show on average a larger normalized burden of meQTLs than inactive TADs (Student-t Test, *p* < 0.05).

### Cancer meQTLs are more abundant in inactive domains

Next we measured the overall burden of independent cancer meQTLs (i.e. meQTLs deemed to represent distinct haplotypes based on the level of linkage disequilibrium; LD) across TAD categories, normalized by TAD length in base pairs. To obtain independent meQTLs, we clumped related meQTLs based on linkage disequilibrium using PLINK. Out of the 1.2 million SNPs, 60,606 remained after LD pruning (**Table 1**). We observed a slightly increased number of cancer meQTLs in inactive domains relative to active regions (Student T-test, p-value < 0.05; **Figure 1C**).

**Table 1:**
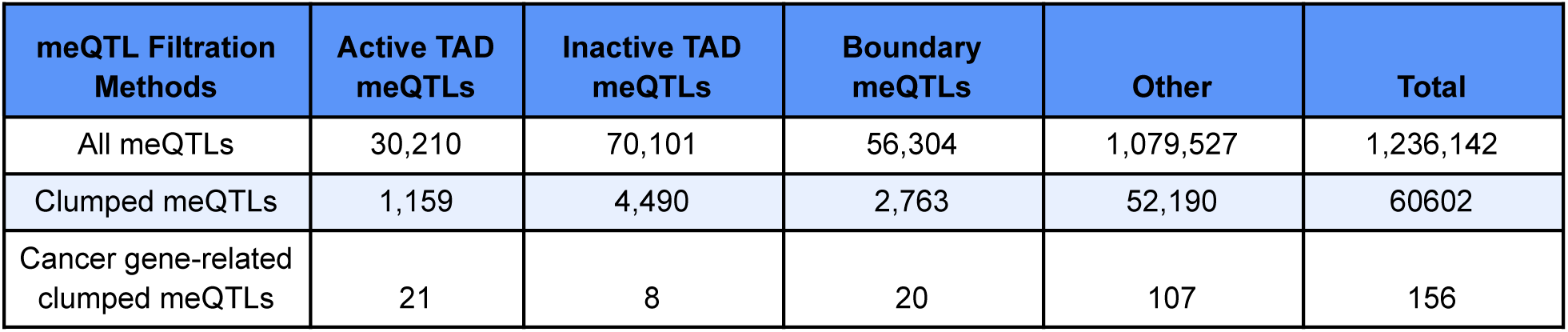
General Information on meQTL Number Across TADs and Multiple Analyses. Each row shows the total number of meQTLs after each analysis across each TAD type. The rows are as follows: all meQTLs without filtration, meQTLs in LD from PLINK clumping software (*p*<1×10^-5^) and meQTLs in LD with CpG probe in cancer driver gene promoter region.

We also evaluated cancer meQTLs at TAD boundaries, considering four categories of boundary based on the category of the flanking TADs: “Active-Boundary-Active”, “Inactive-Boundary-Inactive”, “Active-Boundary-Inactive”, and “Inactive-Boundary-Active”. To allow aggregation across variable length regions, we divided each boundary region into 40 equal genomic bins and calculated the number of meQTLs in each. We then compared the observed density of meQTLs to that obtained by randomizing flanking TAD categories 100 times. Comparing the density of meQTLs in each boundary category to the randomized equivalent, the active-active (student t-test, p<0.01), active-inactive (p<0.01), and inactive-active boundaries (p<0.01) all showed difference in distribution from random, while inactive-inactive (p=0.089) did not (**Figure 2A-D**). Distributions suggested an increase in density of clumped meQTLs when transitioning from active to inactive regions, and conversely, a decrease from inactive to active regions (Kruskal-Wallis ANOVA, p-value<0.05) when compared to the randomly shuffled distribution, but no shift in density for Active-Boundary-Active and Inactive-Boundary-Inactive categories (**Figure 2B-D**).

**Figure 2:**
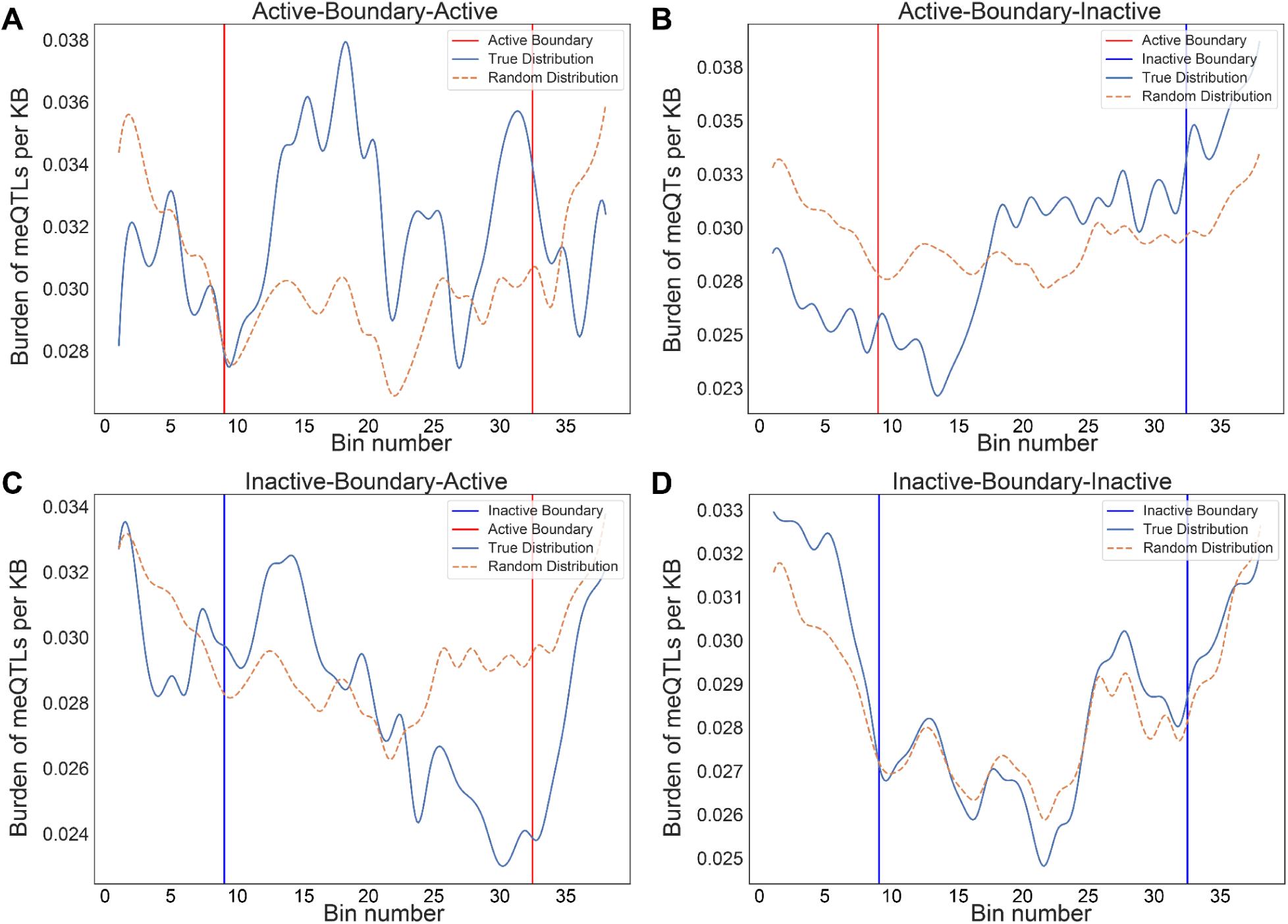
Normalized Burden of meQTLs in Adjacent TADs. The binned average normalized meQTL burden distribution is shown across boundaries between consecutive TADs, grouped by transition category: active to active, active to inactive, inactive to active, and inactive to inactive. The start/end of the TADs for both active and inactive are shown red and blue, respectively. Distributions are smoothened by rolling average for visualization purposes. The graphs represent a unique distribution of meQTL burden across consecutive TADs as opposed to an even spread. The dotted brown line represents the distribution for shuffled random TADs to act as control. (**A**) Active-active (*p=3.51×10^-^*^10^), (**B**) active-inactive (*p=3.45×10^-^*^46^), and (**C**) inactive-active (*p=1.65×10^-^*^25^) boundaries all showed clear difference in distribution from random, while (**D**) inactive-inactive (*p*=0.089) did not.

### Oncogene and tumor suppressor gene-related cancer meQTLs cluster differentially in TADs

Clumped cancer meQTls were further narrowed to those associated with the methylation status of CpG probes located within the promoter regions of cancer driver genes including oncogenes and tumor suppressor genes (TSGs) from the COSMIC database ^16^. In total, 103 oncogenes and 223 TSGs were used for this analysis, where only 67 of them contained meQTL-affecting CpG probes in their promoter regions (i.e. 49 TSGs and 18 oncogenes). Out of the 60,606 clumped meQTLs, 156 of them significantly affected CpG probes located in promoter regions of cancer driver genes (driver meQTLs; **Table 1**). Overall, we saw an overwhelming bias for driver meQTLs to occur in active regions, followed by boundary, and inactive (**Figure 3A**). To understand whether the observed distribution of driver meQTLs was expected, we selected equivalent numbers of meQTLs at random and evaluated their distribution across region types. We did this separately for meQTLs associated with oncogenes versus TSGs, as meQTLs might have different implications in the context of selection for gain versus loss of function. In the oncogene case, meQTLs were depleted relative to random in active TADs, and enriched relative to random in inactive TADs, with no difference in boundary regions. Conversely, for TSGs, there was a significant enrichment of cancer-related meQTLs in active TADs and boundary regions, but a depletion in inactive TADs (**Figure 3B-C**). These opposing trends could suggest genes with the potential to be oncogenes or tumor suppressors (i.e. growth promoting versus limiting) are under different constraints with respect to the propensity for methylation to accumulate in their promoter regions.

**Figure 3:**
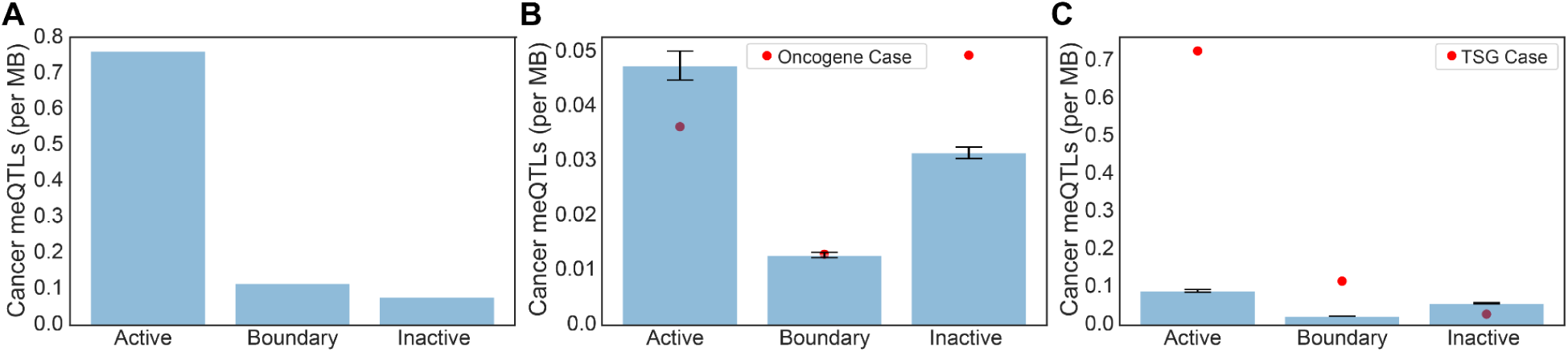
Expected versus Observed Occurrence of Driver meQTLs for Oncogenes and TSGs by Region Type. (**A**) The number of driver meQTLs per MB are plotted, divided according to the category of TAD they are located in. Normalization was conducted by the total region size in each category. (**B-C**) Randomization analysis for burden of non-cancer meQTLs normalized by number of base pairs in each region was conducted to obtain the expected number of cancer meQTLs per MB. To model random expectation **(B**) 54 non-cancer meQTLs (i.e. number of oncogene-proximal meQTLs) and (**C**) 102 non-cancer meQTLs (i.e. number of TSG-proximal meQTLs) were sampled 1000 times for oncogenes and TSGs respectively. Bar graphs are drawn with standard errors. The actual observed cancer meQTL burden is shown as a red dot.

### Assessment of Driver meQTL Association with Cancer Risk and Overall Survival Across Tumor Types

We next evaluated the potential for driver meQTLs to have clinical relevance. A principal component analysis (PCA) was first conducted on the 156 driver meQTLs across individuals in the TCGA. The principal components (PCs) that explained more than 1% of the variance were assessed for association with clinical covariates by linear regression. We noted some association of PCs with tumor type, age at diagnosis and tumor stage at diagnosis, suggesting that cancer meQTLs could have tumor-type specific implications for risk and prognosis. Interestingly, further examining the 10 meQTLs with the strongest loadings in PCs correlated with tumor type, we found that the meQTLs disproportionately affected oncogenes, suggesting that tumor types differ more in oncogene effects than in tumor suppressor effects of DNA methylation.

We first evaluated the driver meQTLs for cancer risk associations using the UKBioBank. In total, 86 of the 155 (1 SNP was not in the UKBB registry) driver meQTLs in the initial PheWAS analysis from UKBioBank patients showed a nominal association with one or more cancer ICD10 codes (p-value<0.05) with 5 SNPs passing a Benjamini-Hochberg FDR threshold of 0.05 (**Table 2**). In total, meQTLs were associated with risk of 15 different cancer types as described by ICD10 codes (**Table 3**). We focused on C50-C50 (malignant neoplasm of the breast) as this tumor type had a large sample size in UKBioBank (n=11,188) and other large cohorts exist to support validation studies.

**Table 2:** List of meQTLs significantly affecting risk and survival in a pan-cancer model (Benjamini-Hochberg FDR<0.05). The beta value is the correlation coefficient of the meQTLs with DNA methylation at the promoter region of the probe gene. The TAD type that the meQTL resides is also represented.

| rsid | SNP | p-value | TAD Type | Probe Gene | Risk/Survival |
| --- | --- | --- | --- | --- | --- |
| rs6500442 | 16:89828862:T:C | 0 | Active | FANCA | Survival |
| rs36083956 | 16:74679883:C:T | 0 | Boundary-Active | RFWD3 | Survival |
| rs1163248 | 10:104896563:A:G | 0 | Neither | NT5C2 | Survival |
| rs1006548 | 16:89844043:T:C | 0 | Active | FANCA | Survival |
| rs8047581 | 16:89884502:C:T | 0 | Active | FANCA | Survival |
| rs17581498 | 17:73794047:G:T | 0 | Inter-TAD | H3F3B | Survival |
| rs62051918 | 16:74613781:T:C | 0 | Active | RFWD3 | Survival |
| rs3935784 | 16:74604841:G:A | 0 | Active | RFWD3 | Survival |
| rs8046036 | 16:74552127:C:T | 0.00000000252 | Active | RFWD3 | Survival |
| rs36030784 | 2:178119204:A:C | 0 | Inter-TAD | NFE2L2 | Survival |
| rs1407920 | 9:10389328:C:G | 0.00000628 | Inter-TAD | PTPRD | Survival |
| rs1725213 | 7:5584599:A:G | 0.00000952 | Active | RAC1 | Survival |
| rs11859725 | 16:74384296:C:T | 0 | Inter-TAD | RFWD3 | Survival |
| rs4265826 | 16:74723707:A:G | 0.0000000519 | Neither | RFWD3 | Survival |
| rs12441344 | 15:67447895:A:G | 0.000000629 | Inter-TAD | SMAD3 | Survival |
| rs200282 | 16:74222799:C:G | 0 | Inter-TAD | RFWD3 | Survival |
| rs6679323 | 1:15914135:A:G | 0 | Active | CASP9 | Survival |
| rs3743861 | 16:89818340:G:C | 0.0000000349 | Active | FANCA | Survival |
| rs10999617 | 10:72723176:G:A | 0.00000886 | Inactive | PRF1 | Risk |
| rs12597188 | 16:68814826:G:A | 0 | Inter-TAD | CDH1 | Risk |
| rs7554885 | 1:18247811:G:T | 0.000000215 | Inter-TAD | SDHB | Risk |
| rs10845664 | 12:13043119:C:T | 0.000000288 | Inter-TAD | CDKN1B | Risk |
| rs741482 | 3:185903412:C:G | 0.0000052 | Inter-TAD | MAP3K13 | Risk |

**Table 3:**
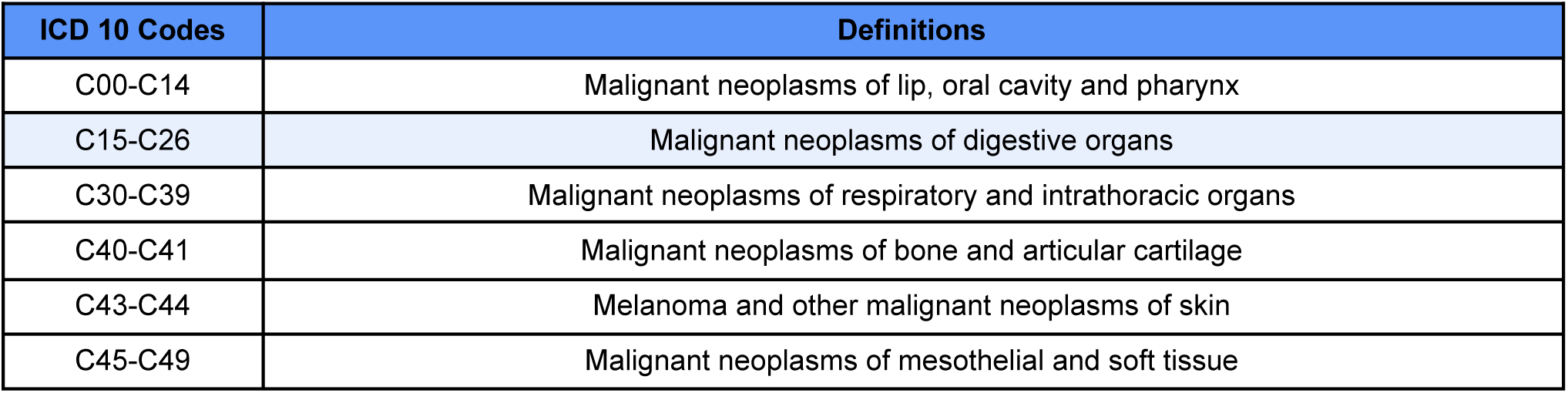
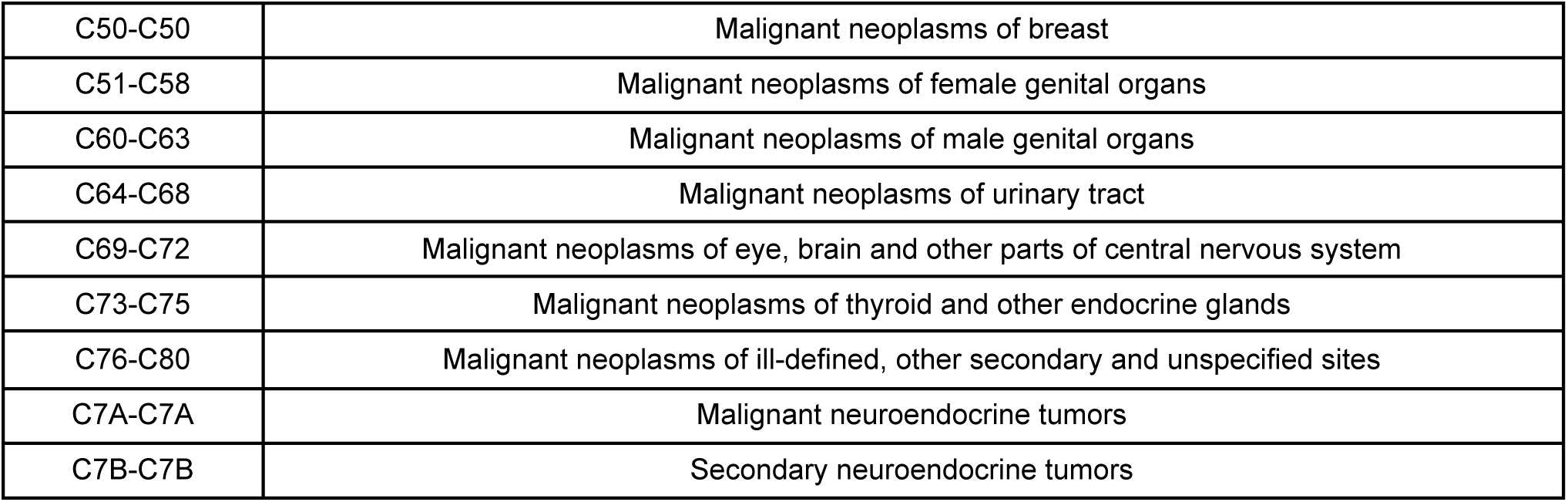
The ICD 10 Code. The ICD 10 code used by UKBB is shown alongside their definitions for the risk analysis.

To further assess the relevance of driver meQTLs to cancer risk, we used them to predict breast cancer status alongside clinical covariates using the approach described by Elgart *et al*^17^. We first performed feature selection by LASSO on nominally significant driver meQTLs and available clinical factors (age, ancestry as represented by the top 10 genotype-derived PCs); LASSO regularization removed ancestry and some meQTLs. Selected features were then used to train an XGBoost classifier on 189,022 examples derived from UKBioBank breast cancer cases and non-cancer controls (**Methods**). The score resulting from the trained XGBoost model was used as the PRS. We applied the trained model to predict breast cancer status for individuals in the DRIVE dataset, comprising 26,374 breast cancer cases and 32,428 controls. The distribution of PRS values across cases was significantly higher than controls for the breast cancer outcome, as expected (Mann-Whitney U, p-value<0.001) (**Figure 4A**). In both UKBB and Drive datasets, the incidence of breast cancer was significantly higher among individuals in the upper 20% percentile of the PRS score versus the bottom 20% percentile (Fisher’s exact test, UKBB: p=4.25×10^-7^<0.001, DRIVE: p=1.47×10^-13^<0.001), suggesting that a higher burden of meQTLs impacts breast cancer risk (**Figure 4B-C**).

**Figure 4:**
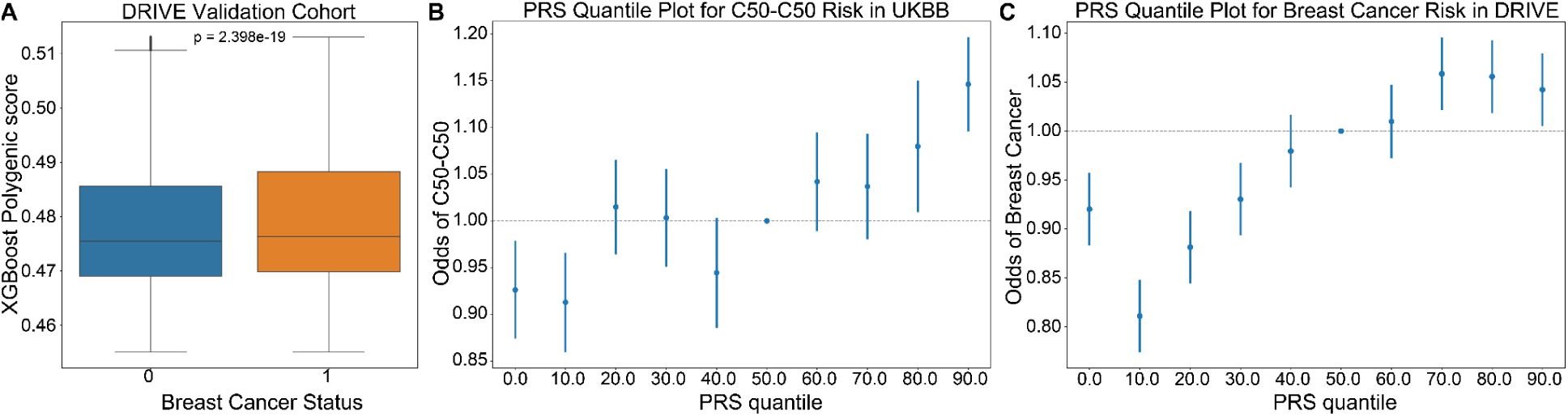
XGBoost Validation of Breast Cancer Risk in DRIVE Dataset. **(A)** An XGBoost classifier trained to predict incidence of breast cancer in the UKBioBank, was applied to predict cancer risk in the DRIVE cohort. PRS scores provided by the model were higher for individuals diagnosed with breast cancer (Figure 8, Mann-Whitney U *p=2.4×10^-^*^19^). (**B-C**) Plots showing the odds ratio of a breast cancer diagnosis across 10% quantiles of the XGBoost predicted PRS in the UKBioBank and DRIVE cohorts respectively. The incidence of breast cancer was significantly higher among individuals in the upper 20% percentile of the PRS score versus the bottom 20% percentile (Fisher’s exact test, UKBB: p=4.25×10^-7^<0.001, DRIVE: p=1.47×10^-13^<0.001) supporting that cancer meQTLs impact breast cancer risk. C50-C50: ICD10 code for malignant neoplasms of the breast.

We extracted feature importances from the UKBioBank-trained PRS to better understand the driver meQTLs underlying breast cancer risk (**Figure 5**). Overall, cancer meQTLs near 29 cancer genes were included in the model. The most predictive driver meQTL was associated MSH2, a gene associated with Lynch syndrome and increased risk of breast cancer ^18^. Polymorphic variation affecting the expression of EZH2, the second most informative feature, has also been linked to breast cancer risk ^19^. ASXL2 may be required for estrogen receptor alpha (ERa) activation in ERa positive breast cancers ^20^. Notably, EZH2 overexpression has been linked more strongly to triple negative breast cancer ^21^ suggesting that the model includes features predictive of multiple subtypes.

**Figure 5.**
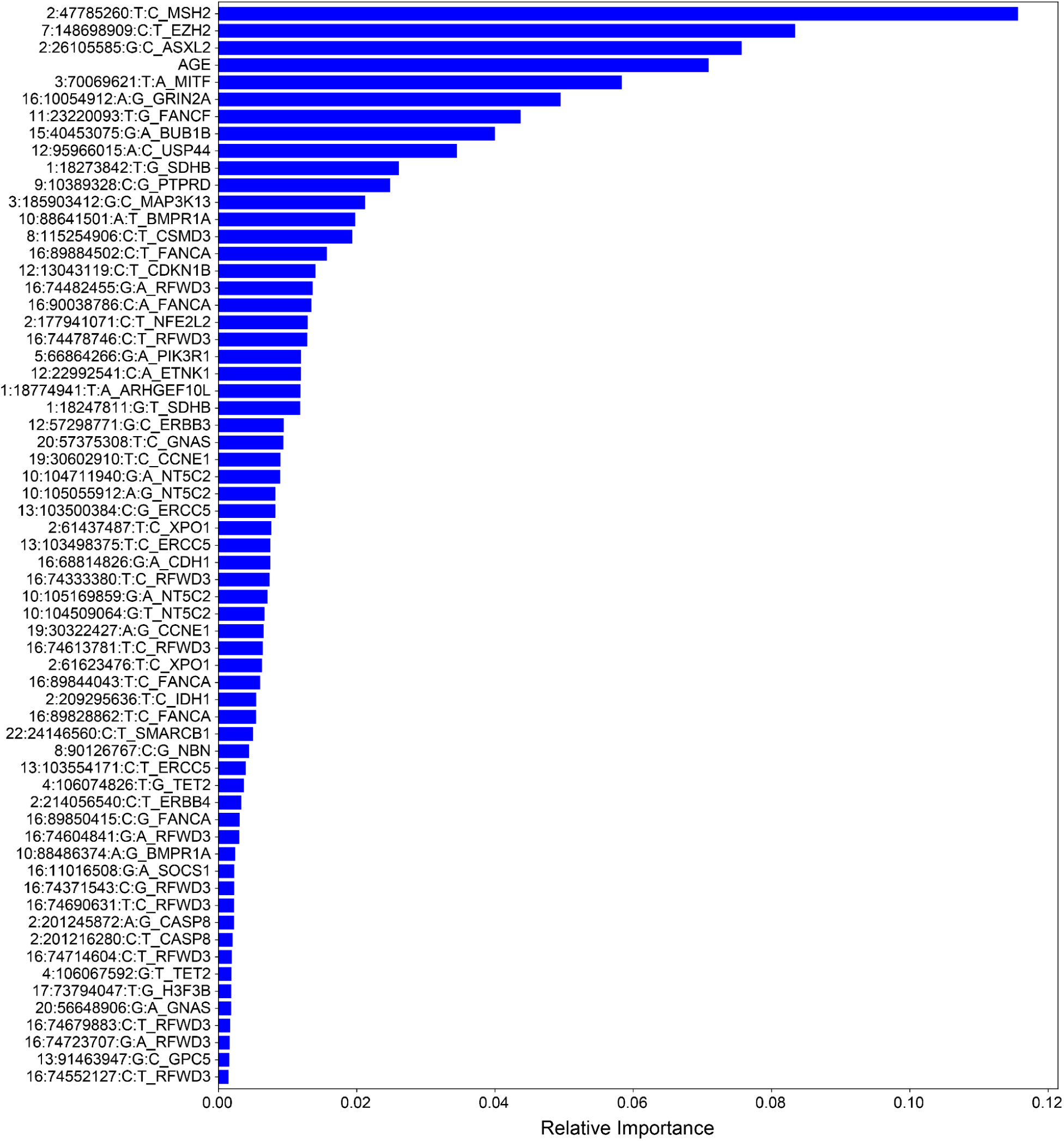
Feature importances for Breast Cancer risk classifier. Features are ranked according to their contribution to classifier predictive performance. Total importances sum to 1.

Finally, we evaluated the implications of driver meQTLs for prognosis. We first removed one meQTL, 2:209220238:C:G, that had a minor allele frequency < 1% across TCGA samples, then conducted a Kaplan-Meier analysis for the remaining meQTLs separately for each tumor type with at least 100 samples. Out of the 155 SNPs, 21 passed the Benjamini-Hochberg adjusted FDR of less than 0.05 (**Table 2**). To assess overall contribution of driver meQTLs to survival, we built polygenic survival scores (PSS) using XGBoost and incorporated them into Cox proportional hazards (PH) models alongside relevant covariates. Here we only evaluated tumor types that had at least 5 SNPs implicated as nominally significant by Kaplan-Meier analysis (n=23 tumor types). Nominally significant driver meQTLs for each tumor type were subjected to selection by LASSO and used to train XGBoost models to predict binary survival outcome (binarized based on median time to an event) separately for each tumor type. Out of the 23 tumor types, 13 had a higher XGBoost classification AUC value when both SNPs and clinical were combined as compared with using only clinical covariates. These included BLCA, BRCA, PAAD, PRAD, UCEC, OV, STAD, SKCM, PCPG, LUSC, KIRC, HNSC and ESCA. This suggests that for these cases, meQTLs contributed survival-relevant information beyond the covariates (*i.e.* age, sex, tumor stage in some cases). For these tumor types, we trained XGBoost models using only meQTLs to obtain tumor-type specific polygenic survival scores (PSS) that were then included alongside covariates (tumor stage, age at diagnosis and sex) in Cox PH models to predict overall survival time in months (**Methods**).

PSS values made a significant contribution to predicting overall survival time for all cancer types except BRCA and SKCM (**Figure 6**). PSS had the highest hazard ratios compared to other covariates for most cancer types, including: ESCA, BLCA, KIRC, LUSC, OV, PAAD, PCPG, PRAD, STAD, UCEC. Most covariates behaved as expected in the analysis with tumor stage having one of the highest odds ratios. However, it is difficult to assess the generalizability of the estimated effect sizes in the absence of independent validation cohorts with both genotype and survival measured in the same cancer types. Nonetheless, to further investigate the prognostic implications of driver meQTLs, we analyzed their feature importances in their respective XGBoost models (**Figure 7**). The number of meQTLs contributing to tumor type specific PSS ranged from 2 to 12, often with 1 or 2 meQTLs dominating the model.

**Figure 6:**
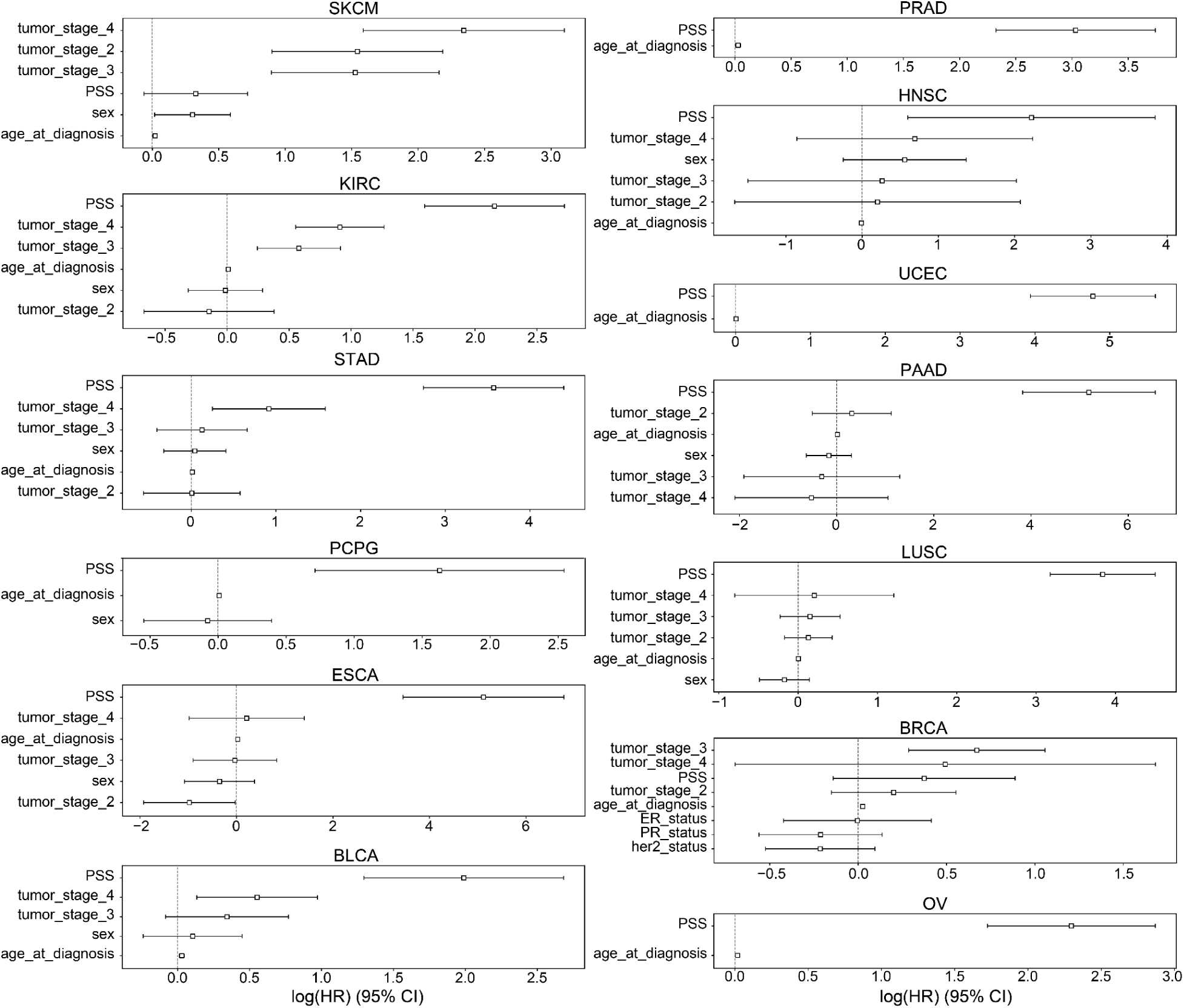
CoxPH 95% Confidence Interval of PSS with Covariates in TCGA Survival. The hazard ratios and 95% confidence intervals associated with various covariates are shown across 13 cancer types: BLCA, BRCA, PAAD, PRAD, UCEC, OV, STAD, SKCM, PCPG, LUSC, KIRC, HNSC, ESCA. Due to limitations in availability of data some tumor types lacked covariates like tumor stage. Sex was excluded for tumors that only occur in males or females. ER: Estrogen receptor, PR: Progesterone Receptor.

**Figure 7:**
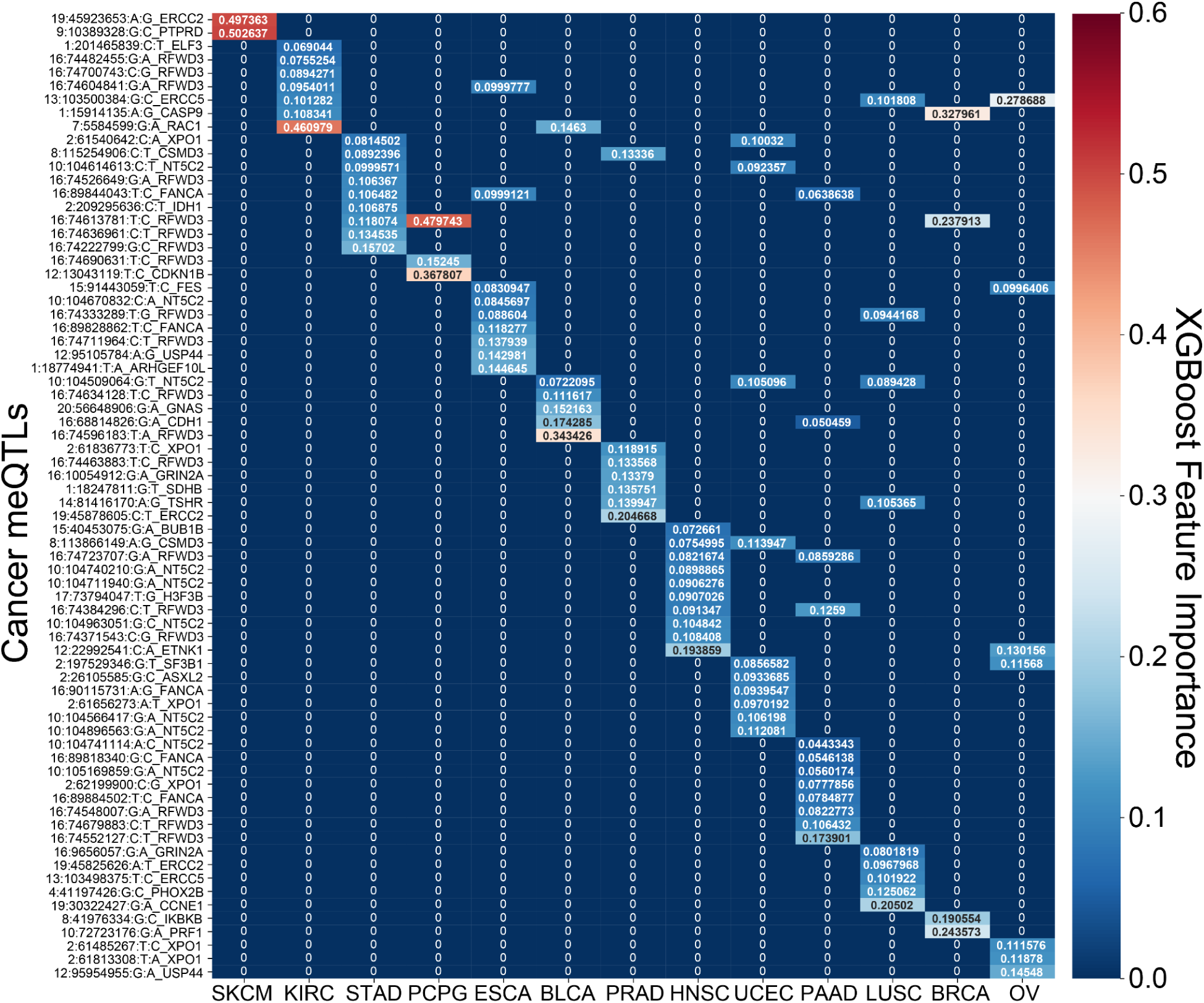
Feature Importance of SNPs in XGBoost Polygenic Survival Scores. A heatmap of the feature importances of SNPs for the cancer type specific XGBoost survival classifiers is shown. For each model across the 13 tumor types, the feature importances sum to 1 with red demonstrating larger importance of a SNP and blue demonstrating lesser importance.

Focusing on the most informative tumor type-associated meQTLs, we investigated the relevance of the associated oncogenes to cancer progression. In many cases, the identified genes were supported by previous studies. For example, PTPRD loss in melanoma was shown to cause disruption of desmosomes, resulting in increased invasive potential ^22^. Polymorphisms in exonuclease ERCC2 have also been found to modify melanoma prognosis ^23^ and have been linked to prostate cancer progression as well ^24^. In pancreatic cancer, RFWD3 expression quantitative trait loci (eQTLs) are associated with survival ^25^. RFWD3 is an E3 protein ubiquitin ligase important for DNA damage and has been shown to stabilize TP53 in response to DNA damage ^26^. We note that RFWD3 meQTLs were among the informative features for many other tumor types as well (**Figure 7**). RAC1 has previously been shown to determine the metastatic potential of renal cell carcinoma (KIRC) ^27^. Reduced expression of CDKN1B is a known risk factor for PCPG, and is common in this disease but usually cannot be explained by somatic alterations, though cases of allelic imbalance have been noted ^28^. CASP9 promoter polymorphisms confer increased risk of breast cancer ^29^ and higher expression of CASP9 was associated with better survival ^30^. Downregulation of ERCC5 is associated with longer progression free survival in ovarian cancer treated with platinum therapy^31^ as is the case for OV in TCGA. In head and neck cancer, the most informative driver meQTL was associated with ETNK1, a cancer gene more commonly associated with myeloid neoplasms^32^ though there is increasing evidence that it may contribute to dysregulation of phospholipid metabolism in multiple tumor types^33^.

## Discussion

There is an increasing appreciation that both genome structure ^34–38^ and common genetic variants ^39–46^ modify to the potential for carcinogenesis. However, the interplay between these factors is not well understood. To start to understand this, we investigated the relationship between the cancer meQTLs recently reported by Gong et al, and 3D genome structure in the form of TADs. To determine the relevance to cancer, we further investigated cancer meQTLs near driver genes for potential to modify cancer risk and progression. We took advantage of a recently introduced modeling strategy that first performs feature selection on a set of nominally associated SNPs, then trains a non-linear XGBoost model based on those features ^17^. Feature importances can be extracted from the trained model to gain insight as to which features were most influential, suggesting biological hypotheses that can be further investigated.

We observed higher levels of promoter methylation in inactive versus active TADs, slightly more meQTLs in active TADs and higher densities of meQTLs in boundary regions proximal to inactive versus active TADs. Furthermore, analyzing meQTL distribution across TAD boundaries revealed a non-uniform pattern, suggesting that TAD boundaries affected distributional burden of meQTLs. It is of note that TAD boundaries conserved across cell types are reportedly highly enriched for evolutionary constraint and complex trait heritability^10^. Interestingly, we found that meQTLs associated with driver genes showed patterns of enrichment or depletion in a manner dependent on the activity state of the TAD in which the meQTLs occurred. Investigating cancer meQTLs, which are polymorphic sites that associate with differences in the level of DNA methylation found in tumors, showed depletion for germline meQTLs affecting oncogenes but enrichment for such meQTLs affecting tumor suppressor genes in active TADs. This could suggest that the potential to modulate tumor suppressor gene expression through methylation is evolutionarily advantageous whereas modulating oncogene expression by promoter methylation may be less so. These trends point to evolutionary constraints on the distribution of meQTLs imposed by 3D genome architectures and that could set the stage for genomic vulnerabilities to later malignancy.

Focusing on meQTLs near known driver genes, we evaluated the potential of meQTLs to modify cancer risk or progression. We found a number of meQTLs associated with survival in the UKBioBank and were able to validate a polygenic score constructed from these meQTLs in the independent DRIVE cohort. The inclusion of genes linked to distinct breast cancer subtypes among the features that most contributed to classifier performance suggests that cancer meQTLs may differentially affect risk of developing different forms of breast cancer and raises the possibility that subtype-specific meQTL-based risk classifiers may outperform a generic model. The meQTLs most strongly predictive of prognosis tended to occur near cancer genes that were also associated with risk or prognosis in the same tumor type. However, we saw cases such as ETNK1 in head and neck cancer, where meQTLs implicated a gene that has not been considered a factor promoting progression. This could point to a new therapeutic opportunity in this disease. Further studies are merited to determine whether the observed associations result from meQTLs being in linkage with eQTLs or coding variants that contribute to risk or progression, or whether meQTLs themselves make it easier or more difficult for genes to be modulated through DNA methylation. Interestingly, we noted multiple independent meQTLs for the same cancer gene were informative in predictive models. This suggests that at least in some cases, the cumulative burden of meQTLs near driver genes could further alter gene function to exacerbate risk or progression. While we focused on cancer genes, others studies have more broadly implicated meQTLs in cancer survival, supporting expanded analyses in the future.

There are a few limitations for this study. First, the meQTLs utilized for this study are derived from a study of tumors ^46^ which could be biased toward detecting meQTLs associated with DNA methylation events that are positively selected in tumors. Second, once focusing on specific tumor types, the number of samples available to predict prognosis is relatively small, and some samples were missing tumor stage or age at diagnosis data, key clinical features for survival prediction. In addition, we lacked independent cohorts to validate the generalizability of polygenic survival scores based on meQTLs, which could lead to overfitting in some of our results as suggested by the large hazard ratios observed in CoxPH analysis. This validation should be a priority as suitable data sets become available. We also made a few assumptions. We only considered common TADs across multiple human cell lines which could have potentially removed some important cell-type specific TAD domains, though our methodology follows what other studies ^11, 47^ have done. For predicting prognosis, we made the assumption that TAD domains from healthy human cell lines would also apply to cancer patients and thus avoided events where TAD structure could change. We justified our decision through previous studies determining TAD domains are overwhelmingly similar across cancer and noncancer patients ^47^. In future studies, it would be of interest to study meQTL trends in normal tissue samples to see if enrichment patterns associated with cancer genes are driven by selection in tumors, or highlight evolutionary constraints more broadly associated with human health that coincidentally are advantageous for tumor development.

This study investigated the relationship between epigenetic factors like chromatin structure and DNA methylation and genetic variation in the context of cancer, and established the potential for survival and risk associated meQTLs to uncover cancer-specific modifiers of risk and progression.

## Methods

### TCGA and Promoter Data

TCGA meQTLs data were obtained from Gong et al ^46^. TCGA outcome and survival data alongside RNA-seq expression data were obtained from the pan-can atlas, Liu et al ^48^. Illumina 450k DNA methylation data were also obtained from the TCGA pan-cancer atlas ^48^. The promoter data was obtained from the ENCODE Screen pipeline ^49, 50^.

### UKBioBank Data

Genotypes and ICD10 codes were obtained for 394,034 samples across 40 ICD 10 codes from the UK BioBank ^51^. For the C50-C50 analysis, only exclusive cases and controls were considered: patients who were only diagnosed with the breast neoplasm were compared with controls who were not diagnosed for any neoplasm. This reduced the sample size to 189,022 for the breast cancer risk analysis.

### DRIVE Breast Cancer Data

Discovery, Biology, and Risk of Inherited Variants in Breast Cancer (DRIVE) (dbGaP Study Accession: phs001265.v1.p1) ^52^ was used to validate the risk outcome analysis of our XGBoost model. There were 60,231 breast cancer cases and controls with genotype data alongside outcome, age, and ancestry principal components.

### TAD Identification and Clustering Based on chromHMM and DNA Methylation

Topologically Associating Domain (TAD) regions from the GM12878, HMEC, HUVEC, IMR90, and NHEK cell lines were downloaded from Rao et al^12^ and only common TAD domains using a 20% overlap algorithm described previously across all 5 cancer cell lines were considered for the rest of the analysis. TAD domains were characterized into 5 clusters: “Active-1”, “Active-2”, “Inactive-1”, “Inactive-2”, and “Mixed” through K-means clustering and use of a 15-chromatin state model derived from the Roadmap Epigenomics Project ^53^. For most of the analysis, the two active and two inactive groups were combined for simpler visualization and mixed regions were ignored due to their biological ambiguity. The boundary of each TAD was considered as the 50 kb region upstream and downstream of TAD endpoints (i.e. 100kb long boundaries) with the exception of consecutive TADs that had a region in between smaller than 100k base pairs. For those cases, the boundary was considered as the proximal half of the region for each of the two TADs. This TAD boundary definition using a 100kb boundary +/-50 kb upstream and downstream from the start and end of a TAD-is supported by previous literature^10^.

DNA methylation levels were compared to TAD domains as follows. DNA methylation levels were summarized at promoters identified by the ENCODE’s SCREEN pipeline for in human hg38. We compared the methylation beta values (i.e. the proportion of methylated region) using TCGA’s DNA methylation data, and averaged these beta values for all promoter regions across Active 1, Active 2, Inactive 1, Inactive 2, and Mixed regions. The hypothesis that methylation levels in promoter regions of actively transcribed TADs would be lower than in inactive TADs was tested by a Kruskal-Wallis test.

### meQTL Distribution within TADs

We retrieved 1,236,142 unique cis-meQTLs across 23 cancer types from the Pancan-meQTL database ^46^. meQTLs were further clumped by Linkage-Disequilibrium (LD) to obtain independent associations using the PLINK ^54^ clumping function using association p-values derived from the Pancan-meQTL database as input and default parameters (p1=0.0001, p2=0.01, r^2^=0.5, kb=250). These clumped, independent meQTLs were used for all subsequent analyses. First, the burden of clumped meQTLs across Active, Inactive, and Mixed TAD regions was measured. The burden was normalized by the length in base pairs of each region. To understand how meQTLs are distributed across the genome and whether TADs have an effect on the distribution of meQTLs, we analyzed the distributional burden of meQTLs within consecutive TADs. We compared the average meQTL density across different TAD transitions(i.e. Active-Boundary-Active, Active-Boundary-Inactive, Inactive-Boundary-Active and Inactive-Boundary-Inactive) by binning the genome between two TADs into 40 equal-sized bins and calculating average burden of meQTLs within these bins normalized by the bin size in base pairs. Resulting graphs were smoothed by a rolling average for visualization purposes. To evaluate whether the distribution reflected an association with transitions in TAD activity status, we shuffled the labels (i.e. “Active”, “Inactive”, etc) of the TADs while preserving the number of transition categories (i.e. “Active-Active”, “Inactive-Active”, etc) 100 times and ran the distribution analysis again on these randomly shuffled TADs by taking an average over all trials. Significance was assessed by comparing the observed difference in density between the TADs to the 100 average randomized trials using a student t-test.

### Randomized Distribution of Cancer-gene-clumped meQTLs

Clumped meQTLs were annotated according to LD with CpG probes located in the promoter regions of cancer driver genes including oncogenes and tumor suppressor genes (TSGs) from the COSMIC database ^16^. A total of 231 oncogenes and TSGs were used for this analysis and promoter regions used were those identified by ENCODE’s SCREEN pipeline ^55^. To evaluate whether active/inactive TADs or boundary regions harboring cancer genes showed enrichment or depletion for meQTLs, we conducted a randomization analysis with 1000 trials. In each trial, we chose a random sample of meQTLs associated with non-cancer genes with matching minor allele frequency (+/-5%) to the set cancer-gene associated meQTLs, while also matching the number of randomly sampled meQTLs. We then mapped genes with meQTLs to active or inactive TADs and TAD boundaries, summed the meQTLs in each and normalized by the size of the region. The standard error was plotted alongside the true burden to see if the burden across TADs is significantly different from random.

### Correlation of meQTL profiles with clinical characteristics in TCGA

We conducted a principal component analysis of TCGA genotype at the 156 meQTLs in European ancestry samples (n=8217), evaluating association of meQTL genotype-based PCs with clinical covariates. meQTL SNPs were quantified by the number of minor alleles carried (0, 1, 2). PCs explaining more than 1% of the genotypic variance across individuals were regressed with clinical variables including sex, age at diagnosis, tumor stage, and tumor type.

### Machine-learning for meQTL-based risk and survival prediction

For both risk and survival analysis, we used a synthesis of LASSO regularization as a feature selector and XGBoost classifier as the machine learning predictor, described fully in Elgart et al ^17^. Specifically, after a preliminary association analysis, SNPs achieving a nominal p-value < 0.05 were further selected by LASSO, and the selected SNPs were used to train an XGBoost model on a predictive task (e.g. cancer versus no cancer for risk, or high survival or low survival at median overall survival time), using a set of training samples. The probabilities achieved from the XGBoost classifier were then used to create a polygenic risk score (PRS) or polygenic survival score (PSS). Predictive performance was evaluated using cross validation for survival analysis and using an independent cohort of matched tumor types for the risk analysis.

### UKBB Risk

To determine the association of meQTLs with risk of developing cancer, we conducted a Phenome-Wide Association Study (PheWAS) for each meQTL using the PLATO ^56^ software. The genotype and phenotype data of 487,409 patients harboring the 156 cancer-related clumped meQTLs was retrieved from the UKBioBank^51^ and genotype at each meQTL was evaluated for association with all cancer phenotypes while controlling for covariates including age and ancestry. Individuals with multiple cancer diagnoses were excluded from the analysis, leaving 189,022 patients for risk analysis.

### UKBB PRS Construction and Breast Cancer Drive Validation

Nominally significant SNPs (p-value<0.05) were used for polygenic risk modeling with LASSO plus XGBoost. Out of the resulting tumor types where meQTLs were associated with risk we pursued breast (ICD-10: C50-C50) due to the abundance of validation data. Of the 189,022 UKBB individuals analyzed, 177,834 and 11,188 patients were non-cancer controls and breast cancer cases, respectively. An initial 10% quantile plot from the Phe-WAS analysis in UKBB was created using the PRS with the odds ratio for C50-C50 to compare the odds ratio of the 0th quantile PRS group to the 90th quantile PRS group.

To create a polygenic risk score (PRS) we utilized the approach described above under “Machine-learning for meQTL-based risk and survival prediction” section. Out of the tumor types that had nominally significant (p<0.05) risk-related SNPs (i.e.C64-C68, C40-C41, C69-C72, C00-C14, C15-C26, C81-C96, C50-C50, C43-C44, C45-C49, C76-C80, C60-C63, C51-C58, C97-C97, C73-C75, C30-C39), we chose to validate this relationship on an external cohort, DRIVE, on the C50-C50 or the breast cancer outcome due to an abundance of validation data. Similar to the survival analysis, we considered SNPs nominally associated with cancer risk using the associations from the PheWAS (p<0.05) for the rest of the analysis. We included other covariates including age and the first 10 principal components to represent population substructure in UKBB. Due to the class imbalance of the UKBB cohort (10,840 cases, 94,871 controls), we oversampled the cases to obtain a 1:1 case control ratio, resulting in a dataset size of 189,742 rows. Furthermore, we only included samples without any neoplasm diagnosis as controls to minimize confounding by other tumor types.

We first trained our XGBoost classification model on the entirety of the UKBB dataset. First the UKBB cohort (i.e. training cohort) was inputted into a LASSO regression model with *α* =0.001 (based on ^17^ ) to predict the intended phenotype. SNPs were further filtered to remove those that had a LASSO coefficient of 0. The modified cohort was used to train an XGBoost model on the filtered feature set using the entire UKBB cohort (n_estimators=500, learning_rate=0.1, max_depth=9). The probability of trees voting for either class (i.e. 0: no cancer, 1:cancer) was used as a polygenic risk score. We validated the breast cancer risk association of meQTLs alongside covariates using the Discovery, Biology, and Risk of Inherited Variants in Breast Cancer (DRIVE ^52^) validation cohort. This validation cohort consists of 32,428 controls and 26,374 breast cancer cases for a total of 58,802 patients. Before validating, we mapped the MAF values of the SNPs in UKBB and DRIVE, and removed SNPs with MAF values of 2 standard deviations away from one another. PRS scores were predicted based on individual genotypes in DRIVE using the UKBB-trained XGBoost model (as described in ^17^). We compared score distributions across case and control in DRIVE using a Mann-Whitney U test. We also compared the incidence of breast cancer by partitioning the UKBB and DRIVE probabilities into 10% quantiles on PRS score. We plotted the 10% quantiles using the min-max normalized XGBoost-derived PRS scores.

### Prediction of survival time in TCGA tumor types

Survival was modeled separately for each of 20 tumor types in TCGA (BLCA, CESC, KIRC, KIRP, PAAD, BRCA, HNSC, LGG, SKCM, PRAD, OV, UCEC, THCA, LUAD, LUSC, COAD, STAD, LIHC, SARC). Cancer meQTLs were included in predictive modeling if they were present with at least 1% minor allele frequency in the specific tumor type, and nominally significant in Kaplan-Meier analysis. Tumor types where fewer than 5 meQTLs showed a nominal association with overall survival or had less than a 100 patients in TCGA were excluded from the analysis. For the remaining tumor types, we divided the analysis into three categories: clinical group containing only clinical features including sex, age, and tumor stage in certain cancer types (i.e. only cancer types >100 patients with non-null tumor stage contained stage as a covariate), control group and SNPs, and SNPs exclusively. For each of the categories, SNPs were selected by LASSO then used the complete dataset to train an XGBoost model, using 5-fold cross validation to estimate the generalization error and generate an AUC value. Specifically, for each individual we simplified the genotypes to a binary feature valued 1 if the patient had the heterozygous or homozygous meQTL allele and 0 if they didn’t. Binarized genotypes were then z-score normalized and input into a LASSO regularization model (*α*=0.001). Features with a LASSO coefficient of 0 (i.e. non-informative features) were removed and the LASSO-filtered SNP set was used to train an XGBoost classifier (n_estimators=500, learning_rate=0.1, max_depth=9) to predict binarized median overall survival (OS, 1=low survival<median survival, 0=high survival>median survival). Cancer types with a higher AUC value in the clinical+SNP group compared to the clinical group were only considered for the SNP only analysis. A higher AUC on the combined group could suggest that SNPs bring additive information. The output of the SNP-only XGBoost model used a non-linear polygenic survival score (PSS). Before inputting into the Cox, the PSS was scaled using the min-max algorithm and outliers were removed using a 1.5*(interquartile range) threshold.

### Cox Proportional Hazard Using PSS

The following tumor types and their corresponding XGBoost regression or PSS scores were used in the Cox model: BLCA, BRCA, CESC, HNSC, KIRC, LUSC, STAD, UCEC. We used Cox proportional hazards models to evaluate the meQTL-based PSS as a predictor of survival interval. We combined the PSS with clinical features including sex, age at diagnosis and tumor stage in a multivariable Cox-proportional hazards model to predict OS, and evaluated the hazard ratios and 95% confidence intervals for each covariate.

## Data and Software Availability

Data were obtained from public sources including The Cancer Genome Atlas, Discovery, Biology, and Risk of Inherited Variants in Breast Cancer (DRIVE) and the UKBioBank. Germline data requires controlled access. meQTLs were obtained from Gong et al ^46^. TADs were obtained from Rao et al^12^. Code to reproduce figures is available at https://github.com/cartercompbio/meQTLs.

## Author Contributions

Original concept by SG and MP. HC supervised the project. SG performed computational data processing and analysis. MP, AK, JT provided support with data set preparation and contributed to computer code. SG, HC wrote the manuscript.

## Acknowledgements

We would like to acknowledge Rany M Salem for providing access to UKBioBank data and TJ Sears for helpful scientific discussion. This research has been conducted using the UK Biobank Resource under Application Number 37671. The results shown here are also based upon data generated by the TCGA Research Network: https://www.cancer.gov/tcga. OncoArray genotyping and phenotype data harmonization for the Discovery, Biology, and Risk of Inherited Variants in Breast Cancer (DRIVE) breast-cancer case control samples was supported by X01 HG007491 and U19 CA148065 and by Cancer Research UK (C1287/A16563). Genotyping was conducted by the Center for Inherited Disease Research (CIDR), Centre for Cancer Genetic Epidemiology, University of Cambridge, and the National Cancer Institute. The following studies contributed germline DNA from breast cancer cases and controls: the Two Sister Study (2SISTER), Breast Oncology Galicia Network (BREOGAN), Copenhagen General Population Study (CGPS), Cancer Prevention Study 2 (CPSII), The European Prospective Investigation into Cancer and Nutrition (EPIC), Melbourne Collaborative Cohort Study (MCCS), Multiethnic Cohort (MEC), Nashville Breast Health Study (NBHS), Nurses Health Study (NHS), Nurses Health Study 2 (NHS2), Polish Breast Cancer Study (PBCS), Prostate Lung Colorectal and Ovarian Cancer Screening Trial (PLCO), Studies of Epidemiology and Risk Factors in Cancer Heredity (SEARCH), The Sister Study (SISTER), Swedish Mammography Cohort (SMC), Women of African Ancestry Breast Cancer Study (WAABCS), Women’s Health Initiative (WHI).

